# Diffusion-MRI-based regional cortical microstructure at birth for predicting neurodevelopmental outcomes of 2-year-olds

**DOI:** 10.1101/2020.04.22.054114

**Authors:** Minhui Ouyang, Qinmu Peng, Tina Jeon, Roy Heyne, Lina Chalak, Hao Huang

## Abstract

Brain cerebral cortical architecture encoding regionally differential dendritic arborization and synaptic formation at birth underlies human behavior emergence at 2 years of age. Brain changes in 0-2 years are most dynamic across lifespan. Effective prediction of future behavior with brain microstructure at birth will reveal structural basis of behavioral emergence in typical development, and identify biomarkers for early detection and tailored intervention in atypical development. Here, we aimed to evaluate the neonate whole-brain cortical microstructure quantified by diffusion MRI for predicting future behavior. We found that individual cognitive and language functions assessed at age of 2 years were robustly predicted by neonate cortical microstructure using support vector regression. Remarkably, cortical regions contributing heavily to the prediction models exhibited distinctive functional selectivity for cognition and language. These findings highlight regional cortical microstructure at birth as potential sensitive biomarker in predicting future neurodevelopmental outcomes and identifying individual risks of brain disorders.

## Introduction

Brain cerebral cortical microstructure underlies neuronal circuit formation and function emergence during brain maturation. Regionally distinctive cortical microstructural architecture profiles around birth result from immensely complicated and spatiotemporally heterogeneous underlying cellular and molecular processes (Silbereis et al., 2016), including neurogenesis, synapse formation, dendritic arborization, axonal growth, pruning and myelination. Disturbance of such precisely regulated maturational events is associated with mental disorders (Innocenti and Price, 2005). Diffusion magnetic resonance imaging (dMRI) has been widely used for quantifying microstructural changes in white matter maturation (e.g. Dubois et al., 2008; Mukherjee et al., 2001). Because of its sensitivity to organized cortical tissue (e.g. radial glial scaffold; Rakic, 1995; Sidman and Rakic, 1973) unique in the fetal and infant brain, dMRI also offers insights into maturation of cortical cytoarchitecture. Cortical fractional anisotropy (FA), a dMRI-derived measurement, of infant and fetal brain can effectively quantify local cortical microstructural architecture and can be used to infer specific brain circuit formation. In early cortical development, most of cortical neurons are generated in the ventricular and subventricular zone. These neurons migrate towards cortical surface along a radially arranged scaffolding of glial cells where relatively high FA values are usually observed (Huang et al., 2013; McKinstry et al., 2002). During emergence of brain circuits, increasing dendritic arborization (Bystron et al., 2008; Sidman and Rakic, 1973), synapses formation (Huttenlocher and Dabholkar, 1997), and myelination of intracortical axons (Yakovlev and Lecours, 1967) disrupt the highly organized radial glia in the immature cortex and result in cortical FA decreases. Such reproducible cortical FA change patterns were documented in many studies of perinatal human brain development (Ball et al., 2013; Huang et al., 2006; Huang et al., 2009; Huang et al., 2013; Kroenke et al., 2007; McKinstry et al., 2002; Neil et al., 1998; Ouyang et al., 2019a; Ouyang et al., 2019b; Yu et al., 2016), suggesting sensitivity of cortical FA measures to maturational processes of cortical microstructure. Diffusion-MRI-based regional cortical microstructure at birth, encoding rich “footage” of regional cellular and molecular processes, may provide novel information regarding typical cortical development and biomarkers for neuropsychiatric disorders.

The first 2 years of life is a critical period for behavioral development, with brain development in this period most rapid across lifespan. In parallel to rapid maturation of cortical architecture and establishment of complex neuronal connections (Hüppi et al., 1998; Ouyang et al., 2019a; Pfefferbaum et al., 1994), babies learn to walk, talk, and build the core capacities for lifetime. Infant behaviors including cognition, language and motor emerge during this time and become measurable at around 2 years of age. Reliable diagnosis for many neuropsychiatric disorders, such as autism spectrum disorder (ASD), can be made only around 2 years of age or later (Marín, 2016), as diagnoses rely on observing behavioral problems which are difficult to recognize in early infancy (Arpi and Ferrari, 2013; Ozonoff et al., 2010). On the other hand, early intervention for ASD, especially before 2 years of age, has demonstrated significant impacts on improving outcomes (Rogers et al., 2014). Given that infants cannot communicate with language or writing in early infancy, there may be no better way to assess their brain development other than neuroimaging. Prediction of future cognition and behavior at 2 years of age or later based on brain features around birth creates an invaluable time window for individualized biomarker detection and early tailored intervention leading to better outcomes.

Individual differences in brain white matter microstructural architectures (Scholz et al., 2009; Yu et al., 2019), behavior, and functions (Braga and Buckner, 2017; Xu et al., 2019) have been well recognized. Individual variability in brain structures and associated individual variability in future behaviors can be harnessed for robust prediction at the single-subject level (Kanai and Rees, 2011; Rosenberg et al., 2018), a step further than group classification. A few studies have been conducted previously to investigate within-sample imaging-outcome correlations (Ball et al., 2015; Counsell et al., 2014; Deoni et al., 2016; Hintz et al., 2015; Keunen et al., 2017; Peyton et al., 2020; Wee et al., 2017; Woodward et al., 2006), while such correlation approaches made it impossible to be applied to new and incoming subjects. Machine learning approach that can adopt new subjects and yield continuous prediction values has been explored only recently based on white matter structural networks (Girault et al., 2019; Kawahara et al., 2017). Cortical microstructure quantified by FA is more directly associated with specific cortical regions and thus certain cortical functions, compared to association of structural networks to the cortical regions through end point connectivity. Our previous study (Ouyang et al., 2019b) demonstrated that cortical FA predicted neonate age with high accuracy. Regionally distinctive cortical microstructure around birth encodes the information that may predict distinctive functions manifested by future behavior and potentially identify the most sensitive regions as imaging markers to detect early behavioral abnormality. However, dMRI-based cortical microstructure has not been evaluated for predicting either discrete or continuous future behavioral measurement so far. And dMRI-based cortical microstructure has not been incorporated into a machine-learning-based prediction model for predicting future behavior, either.

In this study, we leveraged individual variability of cortical microstructure profiles of neonate brains for predicting future behavior. A novel machine-learning-based model using regional cortical microstructure markers from dMRI and capable of predicting continuous outcome values as well as incorporating new subjects was developed. We hypothesized that dMRI-based cortical microstructure at birth only (without inclusion of any white matter microstructure information) could robustly predict the future neurodevelopmental outcomes. Out of 107 recruited neonates, high-resolution (0.656×0.656×1.6 mm^3^) dMRI data were acquired from 87 neonates, of which 46 underwent a follow-up study at their two years of age for neurobehavioral assessments of cognitive, language and motor abilities. Cortical microstructural architectures at birth were quantified by cortical FA on the cortical skeleton to alleviate partial volume effects (Ouyang et al., 2019b; Yu et al., 2016). Regional cortical FA measures were then used to form feature vectors to predict neurodevelopmental outcomes at 2 years of age. We further quantified the contribution of each cortical region in predicting different outcomes, as distinctive behaviors are likely encoded in uniquely distributed pattern across the cerebral cortex.

## Results

### Cortical microstructure at birth and neurodevelopmental outcomes at 2 years of age

A cohort of 107 neonates were recruited for studying normal prenatal and perinatal human brain development (see more details in *Materials and methods* and Supplementary file 1). Neuroimaging data, including structural and diffusion MRI, were collected from 87 infants around birth in their natural sleep. 46 infants went through a follow-up visit at their 2 years of age to complete the cognitive, language and motor assessments with Bayley scales of infant and toddler development-Third Edition (Bayley-III; Bayley, 2006). Figure 1-figure supplement 1 and Figure 1-figure supplement 2 demonstrate the cortical FA maps across parcellated cortical gyri in the left and right hemisphere from dMRI of all these 46 subjects scanned at birth, revealing individual variability of regional cortical microstructure. The Bayley-III composite scores from these 46 subjects at 2 years of age range from 65 to 110 (mean±sd: 87.4±8.5) for cognition, 56 to 112 (85.7±10.1) for language and 73 to 107 (91.2±7.1) for motor abilities (Figure 1-figure supplement 3). No significant correlation between any specific age (i.e. birth age, MRI scan age and Bayley-III exam age) and neurodevelopmental outcome score was found (all *p*>0.1, Supplementary file 2).

### Robust prediction of cognitive and language outcomes based on cortical dMRI measurement

52 cortical regions parcellated by transforming neonate atlas labels (Figure 1, see *Materials and methods*) were used to generate cortical FA feature vectors from each participant’s dMRI data at birth, representing the entire cortical microstructural architecture of an individual neonate. Heterogeneous distribution pattern of cortical FA can be appreciated from cortical FA maps (left panels of Figure 1), indicating regionally differentiated maturation level of cortical microstructure. An immature cerebral cortical region with highly organized radial glia scaffold is associated with high FA values, whereas a more mature cortical region with extensive dendritic arborizations and synapses formations is associated with low FA values. To determine whether cortical microstructural features represented by FA measurements at birth are capable of predicting neurodevelopmental outcomes of an individual infant at a later age, we used support vector regression (SVR) with a fully leave-one-out cross-validated (LOOCV) approach (*middle panels* of Figure 1). With this approach, neurodevelopmental outcome of each infant was predicted from an independent training sample. That is, for each testing subject out of the 46 participants, the cortical FA features of remaining 45 subjects were used to train prediction models for predicting cognitive or language outcomes of the testing subject at 2 years of age (*right panels* of Figure 1) only based on cortical FA of the testing subject at birth. A SVR model that best fits the training sample can be represented by a weighted contribution of all features, where the weight vector 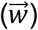 indicates the relative contribution of each feature, namely, cortical FA of each parcellated cortical region, to the prediction model. The feature contribution weights in the model predicting cognition or language were averaged across all leave-one-out SVR models and then normalized. These normalized feature contribution weights were projected back onto the cortical surface to demonstrate cortical regional contribution (*right panels* of Figure 1).

**Figure 1:**
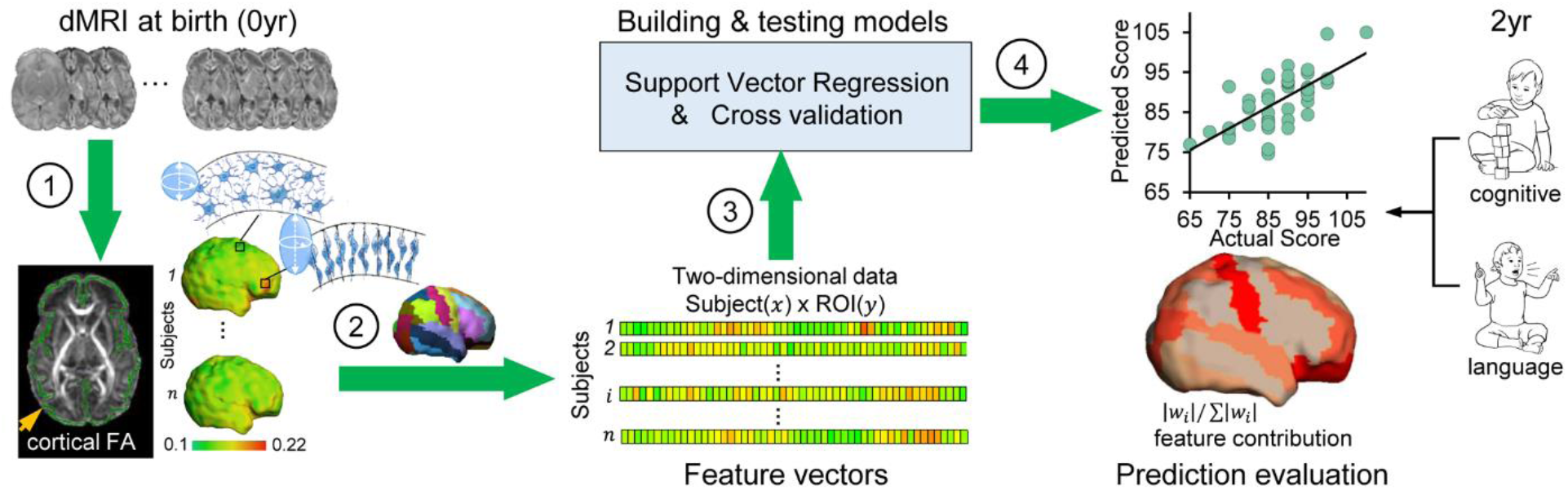
Workflow of predicting neurodevelopmental outcomes at 2 years based on cortical microstructural architecture at birth. Cortical microstructure at birth (0yr) quantified with cortical fractional anisotropy (FA) measures from diffusion MRI (dMRI) was used to predict cognitive and language abilities assessed with Bayley-III Scales at 2 years of age (2yr). The prediction workflow includes the following steps: (1) Cortical microstructure was measured at the “core” of cortical mantle, shown as green skeleton overlaid on a FA map and projected on a neonate cortical surface, to alleviate the partial volume effects. Schematic depiction of dendritic arborization and synaptic formation underlying cortical FA decreases during cortical microstructural maturation is shown. (2) Feature vectors were obtained by measuring cortical skeleton FA at parcellated cortical gyri with the gyral labeling transformed from a neonate atlas. Each parcellated cortical gyrus is a region-of-interests (ROI). (3) Prediction models were established and tested with support vector regression (SVR) and cross-validation. Feature vectors from all subjects were concatenated to obtain the input data of prediction models. (4) Prediction model accuracy was evaluated by correlation between predicted and actual scores. Feature contributions from different gyri in the model were quantified by normalized feature contribution weights which were projected back on a cortical surface for visualization.

Significant correlations between predicted and actual neurodevelopmental outcome were found for both cognitive (*r=0.536, p=1.2*×*10*^*-4*^) and language (*r=0.474, p=8.8*×*10*^*-4*^) scores, respectively (left panel of Figure 2a and Figure 2b), indicating robust prediction of cognitive and language outcomes at 2 years old based on cortical FA measures at birth. According to the permutation tests, these correlations were significantly higher than those obtained by chance (*p*<*0.005*). The mean absolute errors (MAE) between the predicted and actual scores are 5.49 and 7 for cognitive and language outcomes respectively. These MAEs were significantly lower than those obtained by chance (*p*<*0.01*), based on permutation tests. The highly predictive models suggest that cortical microstructural architecture at birth plays an important role in predicting future behavioral and cognitive abilities. However, motor scores were not able to be predicted from cortical FA measures (*r = 0.1, p = 0.52*).

**Figure 2:**
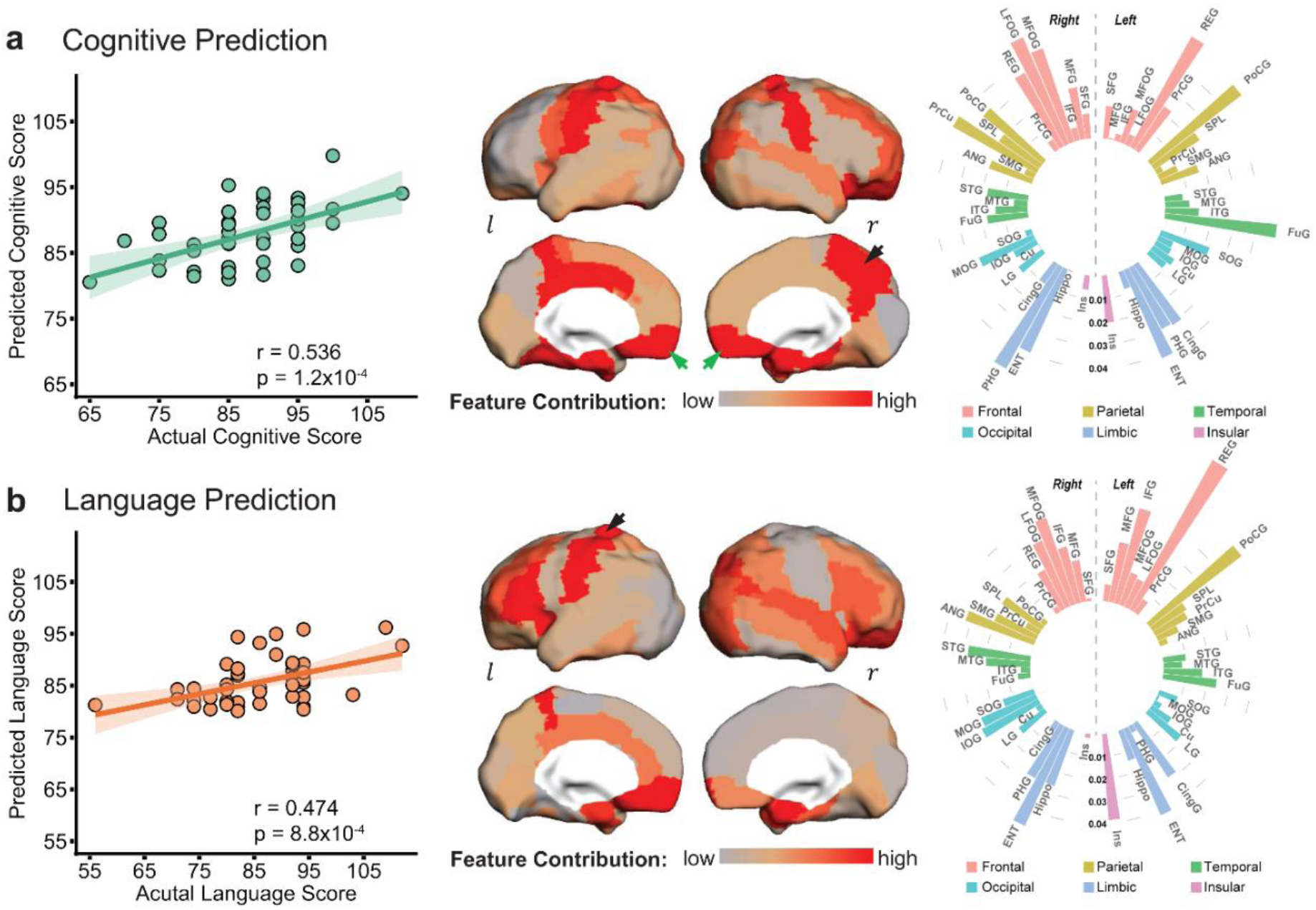
Cortical microstructural measures from neonate dMRI predict cognitive **(a)** and language **(b)** scores at 2 years of age with different feature contribution weights from various cortical gyri. *Left panels:* The scatter plots show significant correlation between actual scores and cognitive (*r=0.536, p=1.2*×*10*^*-4*^) or language (*r=0.474, p=8.8*×*10*^*-4*^) scores predicted based on cortical FA measures. Each dot represents one subject and linear regression was used to assess predictive accuracy of the model. The width of the line denotes the 95% confidence interval around the linear model fit between predicted and observed scores. *Center panels:* Normalized feature contribution weights of all cortical gyri in the prediction models are projected on a cortical surface. *Right panels:* Normalized feature contribution weights from all cortical gyri are demonstrated in the circular bar. These gyri were grouped into frontal, parietal, temporal, occipital, limbic and insular cortex. Abbreviation: *r* : right hemisphere. *l* : left hemisphere. See Supplementary file 3 for abbreviations of cortical regions and values of normalized feature contribution weights from all cortical gyri.

### Evaluation of robustness of prediction models

#### Evaluation with different cortical parcellation schemes and age effects around birth

The prediction models are robust based on evaluation results of different cortical parcellation schemes; and the prediction results are still significant after age adjustment in the cortical FA features (Figure 2-figure supplement 1). To investigate the effects of different cortical parcellation schemes on prediction models, we measured regional cortical FA values with different cortical parcellation schemes that included higher number (128, 256, 512, and 1024) of random cortical parcels. For each parcellation scheme, we calculated correlation coefficient and MAE between the actual and predicted neurodevelopmental scores shown in Figure 2-figure supplement 1. Across different cortical parcellation schemes, robust estimation of the cognitive and language scores was observed in all prediction models. We also investigated the effect of different scan ages on prediction models to demonstrate that high prediction performance remained intact after statistically controlling for age effect in cortical FA measures. Prediction performances before and after adjustment for the age effect are demonstrated in Figure 2-figure supplement 1b-1c. After adjustment for age effect, correlation between the predicted and actual cognitive or language scores is still significant (*p*<*0.05*) with original parcellation of 52 cortical regions. Furthermore, significant correlations after controlling for age effect were also observed across other tested cortical parcellation schemes (128, 256, 512 and 1024 cortical parcels).

#### Evaluation by categorizing subjects with normal and low scores

As Bayley-III is widely used to assess developmental delay with certain cut-off scores, we also evaluated the performance of cortical microstructural measures in classifying subjects with normal and low scores. High accuracy was achieved with a receiver operating characteristic (ROC) curve analysis. Cognitive and language scores of all infants were categorized into normal (>85, n = 22 for cognitive scores and n = 24 for language scores) and low (≤85) scores groups. Cortical FA features were used to build classifiers with leave-one-out procedure to classify each infant into one of these two groups. The ROC curve analysis was used to test the ability of cortical FA measures at birth to distinguish infants with low 2-year-old outcomes from those with normal outcomes (Figure 2-figure supplement 2a). Classification accuracy was 76.1% for cognitive and 60.9% for language scores (Figure 2-figure supplement 2b). Our analysis revealed an area under curve (AUC) of 0.809 and 0.737 for cognitive and language classifications (Figure 2-figure supplement 2c), respectively, further supporting the cortical microstructural architecture at birth as a sensitive marker for prediction and potential detection of early behavioral abnormality.

### Regionally heterogeneous contribution to the cognitive and language prediction

Regional cortical FA measures across entire cortex did not contribute equally to the prediction models. Heterogeneous feature contribution pattern can be clearly seen across cortex for either cognitive (Figure 2a) or language (Figure 2b) prediction. For instance, from distribution of normalized feature contribution weights in cognitive prediction model (*center panel* of Figure 2a), high contributions from right precuneus gyrus (PrCu) (indicated by black arrow) and bilateral rectus gyri (REC) (indicated by green arrows) are prominent, with bright red gyri associated with high feature contribution. To quantitatively demonstrate heterogeneous feature contribution of all cortical gyri, the normalized feature contribution weights from 52 cortical gyri categorized into 6 cortices are shown in a circular bar plot (*right panel* of Figure 2a). Higher bar indicates higher feature contribution of a cortical region to the model. The normalized feature contribution weights of the frontal, parietal and limbic gyri (e.g. REC, postcentral and entorhinal gyri) are relatively higher than those of the occipital, temporal and insular cortex (e.g. superior temporal or occipital gyri) in cognitive prediction.

Similar to cognitive prediction, regional variations of feature contribution can be observed in language prediction model, as demonstrated in cortical surface map and circular bar plot in Figure 2b. For example, higher feature contribution weight was found in the left postcentral gyrus (PoCG) (indicated by black arrow) than its counterpart in the right hemisphere. Feature contribution weights in the frontal and limbic gyri are also higher than those in the occipital and temporal gyri. Differential normalized feature contribution weights in cognitive or language prediction model across all cortical gyri are listed in Supplementary file 3.

### Distinguishable regional contribution to predicting cognitive or language outcomes

Besides regionally heterogeneous contributions, distinguishable feature contribution patterns were found in predicting cognitive or language outcomes. The top 10 cortical regions where microstructural measures contributed most to the prediction of cognitive and language scores are listed in Figure 3a and mapped onto cortical surface in Figure 3b. Among these top 10 cortical regions, left REC, bilateral entorhinal gyrus (ENT), right middle/lateral fronto-orbital gyrus (MFOG/LFOG) and left PoCG are the common regions (highlighted in yellow in Figure 3b) for predicting both cognitive and language outcomes. Right REC, right PrCu, right parahippocampal gyrus (PHG) and left fusiform gyrus (FuG) are unique to cognitive prediction (highlighted in red in Figure 3b), and left inferior frontal gyrus (IFG), left cingular gyrus (CingG), left insular cortex (INS) and right angular gyrus (ANG) are unique to language prediction (highlighted in green in Figure 3b). It is striking that left IFG, usually known as “Broca’s area” for language production, was uniquely found in the top contributing regions in language prediction model. Notably right PrCu, an important hub for default mode network, was uniquely found among the top contributing regions in cognitive prediction model. Bootstrapping analysis indicated that the top 10 cortical regions (Figure 3) where microstructural measures contributing most to prediction were highly reproducible from 1000 bootstrap resamples for predicting each behavioral outcome (Figure 3-figure supplement 1). As shown in Figure 3-figure supplement 1, the cortical regions with higher percentages (indicating higher reproducibility) in red or brown color overlap with the top 10 cortical regions where microstructural measures contributed most to predicting cognition or language (from Figure 3; highlighted by dashed blue contours). Distinguishable regional contribution to predicting different outcomes (cognition or language) was quantified by a nonoverlapping index, ranging from 0 to 1 with 1 indicating completely distinctive regions and 0 indicating same regions.

**Figure 3:**
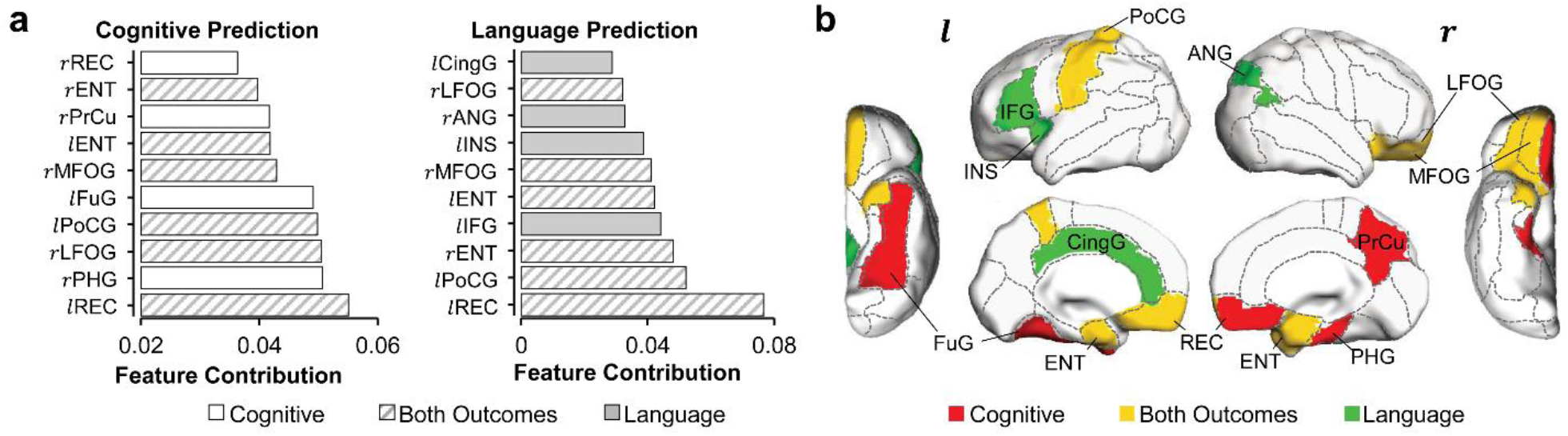
Distinguishable top 10 cortical regions where microstructural measures contributed most to the prediction of cognitive or language scores. **a**, List of top 10 cortical regions with highest feature contribution weights in predicting cognitive (left) or language (right) scores. **b**, Maps of cortical regions listed in **a**. Cortical regions contributing most to predicting both cognition and language are painted in yellow (*r*MFOG, *r* LFOG, *l* REC, *l* PoCG, *r*ENT and *l* ENT); Cortical regions contributing most to predicting uniquely cognition and language are painted in red (*r*REC, *r*PrCu, *l*FuG and *r*PHG) and in green (*l*IFG, *l*CingG, *r*ANG and *l*INS), respectively. *Abbreviation*: *r* : right hemisphere. *l* : left hemisphere. ANG: angular gyrus. CingG: cingular gyrus. ENT: entorhinal gyrus. FuG: fusiform gyrus. IFG: inferior frontal gyrus. INS: insular cortex. LFOG: lateral fronto-orbital gyrus. MFOG: middle fronto-orbital gyrus. PHG: parahippocampal gyrus. PoCG: postcentral gyrus. PrCu: precuneus gyrus. REC: Rectus gyrus.

The statistical significance of the observed nonoverlapping index 0.57 was confirmed with permutation tests. Specifically, the permutation tests indicated that the observed nonoverlapping index of 0.57 was not likely to be obtained by chance from predicting the same outcome (p=0.001 from testing with leave-one-out resamples; p=0.05 from testing with resamples by randomly selecting 90% of samples), supporting distinguishable regional contribution to predicting cognitive or language outcomes.

### Comparison between prediction based on cortical FA and prediction based on white matter FA

Since dMRI has been conventionally used mainly for measuring white matter microstructure, we also evaluated the prediction performance when only using regional white matter FA measures as features (Figure 4a). Both cognitive (*r=0.516, p=2.4*×*10*^*-4*^) and language (*r=0.517, p=2.3*×*10*^*-4*^) scores can be reliably predicted with white matter FA measures, indicated by significant correlations between predicted and actual scores (Figure 4b). More importantly, solely cortical FA measures at birth is as robust as white matter FA measures in predicting the cognitive and language scores at 2 years of age, demonstrated by similar correlation coefficient values between the predicted and actual scores (Figure 4c, *all* significant correlation with *p*<*0.05*).

**Figure 4:**
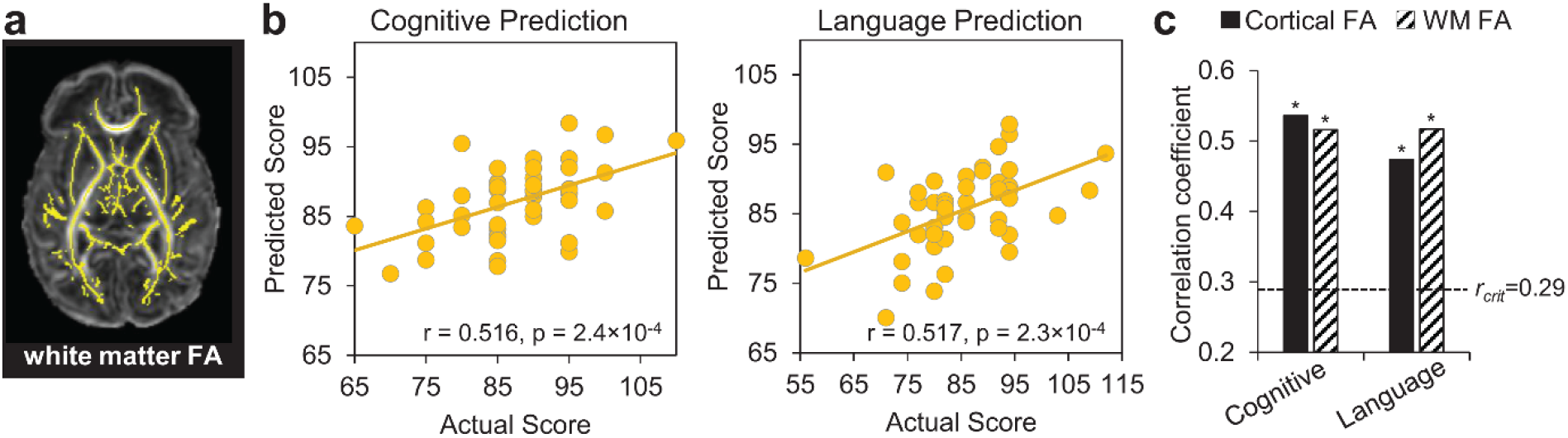
Prediction of neurodevelopmental outcomes using white matter (WM) FA features, compared with prediction using cortical FA features. **a**, WM microstructure was measured at the core WM regions, shown as yellow skeleton overlaid on a FA map of a neonate brain, to alleviate the partial volume effects. Feature vectors were obtained by measuring WM skeleton FA at 40 tracts transformed from WM labeling of a neonate atlas. **b**, Scatter plots show linear regressions between actual scores and the predicted cognitive or language scores based on WM FA measures with LOOCV. **c**, Significant correlation between the predicted and actual cognitive or language outcomes was found based on both cortical FA and WM FA feature vectors. Dashed line indicates critical *r* value corresponding to *p=0.05*. * in the panel indicates significant (*p*<*0.05*) correlation.

## Discussion

We leveraged individual variability in the cortical microstructural architecture at birth for a robust prediction of individualized future behavioral outcomes in continuous values. Cortical microstructure at birth, encoding rich “footage” of regional cellular and molecular processes in early human brain development, was evaluated as the baseline measurements for prediction. Our previous studies (Huang et al., 2009; Ouyang et al., 2019b; Yu et al., 2016) found that individual neonate cortical microstructure profile characterized by different levels of dendritic arborization could be reliably quantified with dMRI-based cortical FA that reveals cortical maturation signature. In this study, we further demonstrated that individual variability of cortical microstructure profile around birth can be used to robustly predict the cognitive and language outcomes of individual infant at 2 years of age. By harnessing different encoding patterns of cognitive and language functions across the entire cortex, we quantified distinguishable contribution of each cortical region and highlighted the most sensitive regions for predicting different outcomes. Cortical regions contributing heavily to the prediction models exhibited distinctive functional selectivity for cognition and language. To our knowledge, this is the first study evaluating regional cortical microstructure for predicting future behavior, laying the foundation for future works using cortical microstructure profile as ‘neuromarkers’ to predict the risk of an individual developing health-related behavioral abnormalities (e.g. ASD). The prediction model is also capable of incorporating new and incoming subjects, a step further than previous within-sample imaging-outcome correlation studies (Ball et al., 2015; Counsell et al., 2014; Deoni et al., 2016; Hintz et al., 2015; Keunen et al., 2017; Peyton et al., 2020; Wee et al., 2017; Woodward et al., 2006). If given a larger sample size, this study has the potential to change the paradigm of identifying infants at risk of neurodevelopmental disorders (e.g. ASD) at a time when infant is pre-symptomatic in behavioral assessments and intervention can lead to better outcomes.

With cerebral cortex playing a central role in human cognition and behaviors, high performance achieved in prediction of cognition and language based on cortical FA measures (Figure 2) is probably due to sensitivity of cortical microstructural changes to maturational processes involving synaptic formation, dendritic arborization and axonal growth. Distinctive maturation processes manifested by differentiated cortical FA changes across cortical regions in fetal and infant brains were reproducibly reported development (Ball et al., 2013; Huang et al., 2006; Huang et al., 2009; Huang et al., 2013; Kroenke et al., 2007; McKinstry et al., 2002; Neil et al., 1998; Ouyang et al., 2019a; Ouyang et al., 2019b; Yu et al., 2016). Although cortical thickness, volume or surface area from structural MRI scans (i.e. T1- or T2-weighted images) were conventionally primary structural measurements to characterize infant cerebral cortex development (Hazlett et al., 2017; Hill et al., 2010; Lyall et al., 2015), they cannot characterize the complex microstructural processes that take place inside the cortical mantle. Compared to macrostructural changes quantified by these conventional measurements, the underlying microstructural processes quantified by cortical FA may be more sensitive to infants with pathology such as those with risk of ASD. Because white matter microstructure (e.g. FA) measurement is more widely used in dMRI studies than cortical microstructure measurement, we also evaluated white matter FA at birth for predicting cognitive and language outcome at 2 years of age (Figure 4). Despite the fact that microstructural measures from solely cerebral cortex or white matter at birth have similar sensitivity in predicting neurodevelopment outcomes at 2 years old (Figure 4), cortical microstructure is more directly associated with specific cortical regions and thus certain cortical functions, compared to association of white matter to the cortical regions through end point connectivity.

Human brain development in the first two years is most rapid across the lifespan. During first 2 years after birth, overall size of an infant brain increases dramatically, reaching close to 90% of an adult brain volume by 2 years of age (Pfefferbaum et al., 1994). Despite rapid development, infancy period (0-2 years) of human is probably the longest among all mammals with cognitive and language functions unique in human emerging during this critical period. For instance, infants start to learn their mother tongue from babbling to full sentences, during age of 6-month to 2-3 years (Kuhl, 2004). This lengthy yet extremely dynamic brain development processes make the prediction of 2-year neurodevelopmental outcome with brain information at birth ultimately invaluable. DMRI-based cortical microstructure at birth, before any behavioral tests could be performed, can well predict both cognition and language outcomes at 2 years of age (Figure 2). A regionally heterogeneous distribution pattern across cerebral cortex was displayed (Figure 2). Furthermore, consistent to their functions documented in the literature, the cortical regions contributing heavily to the prediction models exhibited distinguishable functional selectivity for cognition and language (Figure 3 and Figure 3-figure supplement 1). The cognitive scale in Bayley-III estimates cognitive functions including object relatedness, memory, problem solving and manipulation on the basis of nonverbal activities (Bayley, 2006). Cortical regions with high weight in prediction model are tightly associated with cognitive functions. Right rectus, precuneus, parahippocampal and left fusiform gyri were among the top 10 cortical regions (painted red in Figure 3b) where microstructural measures contributed uniquely to the cognition prediction, but not to language prediction. Parahippocampal gyrus provides poly-sensory input to the hippocampus (Witter et al., 2000) and holds an essential position for mediating memory function (Young et al., 1997). Precuneus is a pivotal hub essential in brain’s default-mode network (Buckner et al., 2008) and involved in various higher-order cognitive functions (Cavanna and Trimble, 2006). Rectus and fusiform gyri are associated with higher-level social cognition processes (Viskontas et al., 2007). For regions contributing heavily to both cognition and language prediction and painted yellow in Figure 3b, bilateral entorhinal gyri likely serve as a pivot junctional region mediating the processes of different types of sensory information during the cortex-hippocampus interplay (Witter et al., 2000). Middle and lateral fronto-orbital gyrus are involved in cognitive processes including learning, memory and decision-making (Wikenheiser and Schoenbaum, 2016). Left postcentral gyrus with high feature contribution weight in both prediction models is associated with the general motor demands of performing tasks. Besides higher-order cognitive functions, language functions are also unique in human beings. The first two years of life is a critical and sensitive period for the speech-perception and speech-production development (Kuhl, 2004; Werker and Hensch, 2015). The language scale from Bayley-III includes two subdomains: receptive communication and expressive communication. Here, the term “communication” refers to any way that a child uses to interact with others, and includes communication in prelinguistic stage (e.g. eye gaze, gesture, facial expression, vocalizations, words), social-emotional skills and communication in more advanced stage of language emergence (Bayley, 2006). The distinctive regions with high feature contribution weights only in language prediction model included left inferior frontal gyrus (IFG), cingular gyrus, insular cortex and right angular gyrus, painted green in Figure 3b. These regions are related to receptive and expressive communication. It is striking that left IFG, known as “Broca’s area”, was identified by this data-driven prediction model because it is well known that Broca’s area plays a pivotal role in producing language (Poeppel, 2014). Angular gyrus, another crucial language region in the parietal lobe, supports the integration of semantic information into context and transfers visually perceived words to Wernicke’s area. Left insula, a part of the articulatory network in the dual-stream model of speech processing, is involved in translating acoustic speech signals into articulatory representations in the frontal lobe (Hickok and Poeppel, 2007). Since Bayley language scale also includes a number of items reflecting social-emotional skills, such as how a child responds to his/her name or reacts when interrupted in play, high feature contribution weight of insular cortex may be due to its important role in social emotions (Lamm and Singer, 2010). High feature contribution weight of cingular cortex might be related to its key involvement in emotion and social behavior (Bush et al., 2000). Taken together, identifying these regions with highest feature contribution weights sheds light on understanding local brain structural basis underlying emergence of distinctive functions manifested by daily behavior, enhancing our knowledge of brain-behavior relationships.

Motor scores from Bayley-III were not predicted reliably in this study possibly due to low variability and low signal-to-noise ratio of cortical FA measurement in primary sensorimotor cortical regions associated with motor function. Cortical FA measurements at primary sensorimotor cortex are relatively low compared to those at other cortices (Ball et al., 2013; Ouyang et al., 2019b) and are barely above the noise floor, as primary sensorimotor cortex develops earlier compared to cortical regions associated with higher-order brain functions. Individual variability of cortical microstructure at primary sensorimotor cortex cannot be well captured with relatively low signal-to-noise ratio for the cortical FA measurements at these regions. High individual variability enables reliable prediction. Low functional variability at primary sensorimotor cortex was found in a largely overlapped cohort in a separate study (Xu et al., 2019) from our group, and was reproducibly found in another cohort (http://developingconnectome.org). With strict exclusion criteria of participating cohort around birth, higher average and lower variance of motor scores than those of cognition or language scores play an important role in poor prediction of motor scores. Larger group variability in motor scores and larger sample size can offset the limitations elaborated above and enhance the prediction of motor scores.

Technical considerations, limitations and future directions are discussed below. The capacity of cortical microstructural profile at birth to predict an individual infant’s later behavior is substantial (Figure 2). We trained models to classify the low and normal outcomes. The ROC curves and accuracy measurements in Figure 2-figure supplement 2 demonstrated high accuracy of the classification models. Robustness of the prediction model was further tested against various factors. These factors included different cortical parcellation schemes (random and finer parcellations versus parcellation based on an atlas) for measuring feature vectors, and individual age adjustment (Figure 2-figure supplement 1). Importantly, high performance of the prediction models is reproducible after taking above-mentioned factors into consideration. Despite relatively high dMRI resolution (0.656×0.656×1.6 mm^3^) being used, partial volume effects (Jeon et al., 2012) cannot be ignored for measuring cortical FA. The partial volume effects are different across brain with thinner cortical regions more severely affected. To maximally alleviate the partial volume effects and enhance the measurement accuracy, we adopted a “cortical skeleton” approach (Ouyang et al., 2019b; Yu et al., 2016), demonstrated in the left panel of Figure 1, to measure cortical microstructure at the center or “core” of the cortical plate. Although both preterm and term-born infants are included, none of them were clinically referred. All infants had been recruited solely for brain research and rigorously screened by a neonatologist and a pediatric neuroradiologist to exclude any infants with signs of brain injury (see *Materials and method* for more details). To limit the effects of exposure to the extrauterine environment, this study was designed to make the interval between birth and scan age as short as possible. As a result, we did not find any significant correlation between birth age or MRI scan age and neurodevelopmental outcomes (see Supplementary file 2). The literature (Bonifacio et al., 2010) also indicated that the effects of premature birth on brain development are considered to be relatively trivial compared with the effects of brain injury and co-morbid condition which was not presented in any recruited infant due to strict exclusion criteria. Although we have taken many precautions to extract cortical FA measures and test internal validity of our prediction analysis, several limitations will need to be addressed in future research. Despite the fact that relatively high performance of behavioral prediction was achieved with current cohort of infants, the prediction model will benefit from validation (*e.g. k*-fold cross-validation) and replication with an independent infant cohort of a larger sample size for generalization. Thus, prediction model with individual variability representing a general population from a much larger cohort is warranted in future research. Future research will also benefit from incorporating other markers such as functional neuroimaging, multimodal imaging and genetic factors (*e.g.* Kwon et al., 2014; Smyser et al., 2016) as well as adopting more advanced machine learning algorithms such as multi-kernel and deep learning to further improve prediction. This study included a healthy cohort of infants for evaluating cortical microstructure for predicting future behavior. Such evaluation in the setting of pathology needs to be further validated. The observed neurodevelopmental outcomes were also contributed by unmeasured factors such as maternal age and tertiary educational level as well as other home environment variable following discharge from the hospital, all of which should be taken into consideration in the future prediction model.

In conclusion, whole-brain cortical FA at birth, encoding rich information of dendritic arborization and synaptic formation, could be reliably used for predicting neurodevelopmental outcomes of 2-years-old infants by leveraging individual variability of these measures. Feature contribution weight in cognitive or language prediction is heterogeneous across brain regions. The cortical regions contributing heavily to the prediction models exhibited distinguishable functional selectivity for cognition and language. Identifying regions with highest feature contribution weights offers preliminary findings on understanding local brain microstructural basis underlying emergence of future behavior, enhancing our knowledge of brain-behavior relationships. These findings also suggest that cortical microstructural information at birth may be potentially used for prediction of behavioral abnormality in infants with high risk for brain disorders early at a time when infant is pre-symptomatic in behavioral assessments and intervention may be most effective.

## Materials and methods

### Participants

The study was approved by the Institutional Review Board (IRB) at the University of Texas Southwestern Medical Center. 107 neonates were recruited from the Parkland Hospital and scanned at Children’s Medical Center at Dallas. Evaluable MRI was obtained from 87 neonates (58 M/ 29 F; post-menstrual ages at scan: 31.9 to 41.7 postmenstrual weeks (PMW); post-menstrual ages at birth: 26 to 41.4 PMW). All recruited infants were not clinically indicated, and they were recruited completely for research purpose which was studying the normal prenatal and perinatal human brain development. These neonates were selected through rigorous screening procedures by a board-certified neonatologist (LC) and an experienced pediatric radiologist, based on subjects’ ultrasound, clinical MRI and medical record of the subjects and mothers. The exclusion criteria included evidence of bleeding or intracranial abnormality by serial sonography; the mother’s excessive drug or alcohol abuse during pregnancy; periventricular leukomalacia; hypoxic–ischemic encephalopathy; Grade III–IV intraventricular hemorrhage; body or heart malformations; chromosomal abnormalities, lung disease or bronchopulmonary dysplasia; necrotizing enterocolitis requiring intestinal resection or complex feeding/nutritional disorders; defects or anomalies of the brain; brain tissue dysplasia or hypoplasia; abnormal meninges; alterations in the pial or ventricular surface; or white matter lesions. Informed parental consents were obtained from the subject’s parent. More demographic information of the participants can be found in Supplementary file 1.

### Neonate brain MRI

All neonates were scanned with a 3T Philips Achieva System (ages at scan: 31.9 to 41.7 PMW). Neonates were fed before the MRI scan and wrapped with a vacuum immobilizer to minimize motion. During scan, all neonates were asleep naturally without sedation. Earplugs, earphones and extra foam padding were applied to reduce the sound of the scanner. All 87 neonates underwent high-resolution diffusion MRI (dMRI) and structural MRI scans. A single-shot echo-planar imaging (EPI) sequence with Sensitivity Encoding parallel imaging (SENSE factor = 2.5) was used for dMRI. Other dMRI imaging parameters were as follows: time of repetition (TR) = 6850ms, echo time (TE) = 78ms, in-plane field of view = 168 ×168 mm^2^, in-plane imaging matrix = 112 ×112 reconstructed to 256×256 with zero filling, in-plane resolution = 0.656 ×0.656 mm^2^ (nominal imaging resolution 1.5 ×1.5 mm^2^), slice thickness = 1.6mm without gap, slice number = 60, and 30 independent diffusion encoding directions with b value = 1000s/ mm^2^. Two repetitions were conducted for dMRI acquisition to improve the signal to noise ratio (SNR), resulting in a scan time of 11 minutes.

### Quality control and quality assurance of MRI

General MRI slice and slice-time integral measures for quality control (QC) were determined daily using ADNI and BIRN phantoms. Any systematic anomaly identified by significant deflections from normal variation was addressed immediately with technical support and/or the in-house MR physicist team. As is the laboratory practice, test-retest reliability of the MR imaging protocol was assessed with a 4 subject X 4 repeat estimation on intra- and inter-subject variation for quality assurance (QA).

### Measurement of cortical microstructure with brain MRI at birth

Diffusion tensor of each brain voxel was calculated with routine tensor fitting procedures. With 30 scanned diffusion weighted image (DWI) volumes and 2 repetitions, we accepted those scanned diffusion MRI datasets with less than 5 DWI volumes affected by motion more commonly seen in scanning of neonates and toddlers. Diffusion MRI datasets from all neonates were preprocessed using *DTIstudio* (http://www.mristudio.org) (Jiang et al., 2006). Small motion and eddy current of dMRI for each neonate were corrected by registering all the DWIs to the non-diffusion weighted b0 image using a 12-parameter (affine) linear image registration with automated image registration algorithm. Six elements of diffusion tensor were fitted in each voxel. Maps of fractional anisotropy (FA) derived from diffusion tensor were obtained for all neonates (Figure 1). DTI-derived FA maps were used to obtain the cortical skeleton FA measurements at specific cortical gyral region of interests (ROI) identified by certain gyral label from a neonate atlas (Feng et al., 2019). To alleviate partial volume effects, the cortical FA values were measured on the cortical skeleton, i.e. the center of the cortical mantle, demonstrated as green skeletons in the left panels of Figure 1. This procedure was elaborated in our previous studies (Ouyang et al., 2019b; Yu et al., 2016). The cortical skeleton was created from averaged FA maps in three age-specific templates at 33, 36 and 39PMW due to dramatic anatomical changes of the neonate brain from 31.9 to 41.7 PMW. Based on the scan age, individual subject brain was categorized into 3 age groups at 33, 36 and 39PMW, and registered to the corresponding templates using the registration protocol described in details in the literature (Feng et al., 2019; Oishi et al., 2011). By applying the skeletonization function in *TBSS* of *FSL* (http://fsl.fmrib.ox.ac.uk/fsl/fslwiki/TBSS), cortical skeleton of the 33PMW or 36PMW brain was extracted from the averaged cortical FA map and cortical skeleton of the 39PMW brain was obtained with averaged cortical segmentation map due to low cortical FA in 39PMW brains. The cortical skeleton in the 33, 36 and 39PMW space was then inversely transferred to each subject’s native space, to which the 52 cortical gyral labels of a neonate atlas (Feng et al., 2019) were also mapped to parcellate the cortex (Figure 1). By directly overlapping the cortical skeleton with the neonate atlas, the cortical skeleton was parcellated into 52 gyri. The FA measurement at each cortical gyrus was calculated by averaging the measurements on the cortical skeleton voxels with this cortical label. In this way, feature vectors consisting cortical FA values from 52 parcellated cortical gyri and measured at the cortical skeleton were obtained for the following support vector regression (SVR) procedures.

### Neurodevelopmental assessments at 2 years of age

Out of 87 neonates with evaluable MRI scanned around birth, a follow-up neurodevelopmental assessment was obtained from 46 neonates (32M/14F, scan age of 36.7±2.8PMW) at their 2 years of age (20-29months, 23.5±2.3months) corrected for prematurity, with gestational age taken into account. Cognitive, language and motor development were assessed using the Bayley-III (Bayley, 2006). Specifically, the cognitive scale estimates general cognitive functioning on the basis of nonverbal activities (i.e. object relatedness, memory, problem solving, and manipulation); the language scale estimates receptive communication (i.e. verbal understanding and concept development) as well as expressive communication including the ability to communicate through words and gestures; and the motor scale estimates both fine motor (i.e. grasping, perceptual-motor integration, motor planning and speed) and gross motor (i.e. sitting, standing, locomotion and balance) (Bayley, 2006). The Bayley-III is age standardized and widely used in both research and clinical settings. It has published norms with a mean (standard deviation) of 100 (15), with higher scores indicating better performance. This neurodevelopmental assessment was conducted by a certified neurodevelopmental psychologist, who was blinded to clinical details of infants as well as the neonate MR findings. Unlike cognitive, language and motor scales reliably obtained using items administered to the child by a certified neurodevelopmental psychologist, other two scales from Bayley-III (social-emotional and adaptive scales) obtained from primary caregiver heterogeneous responses to questionnaires were not included in this study.

### Prediction of neurodevelopmental outcome with cortical FA as features

To determine whether cortical FA at birth could serve as a biomarker for individualized prediction of neurodevelopmental outcomes at 2 years of age, we performed pattern analysis using SVR algorithm implemented in *LIBSVM* (Chang and Lin, 2011). SVR is a supervised learning technique based on the concept of support vector machine (SVM) to predict continuous variables such as cognitive, language or motor composite score from Bayley-III. Leave-one-out cross-validation (LOOCV) was adopted to evaluate performance of the SVR model for each score. Cortical FA at birth from one individual subject was used as the testing data and the information of remaining 45 subjects including their cortical FA at birth and Bayley scores at 2 years of age were used as training data. In this procedure, the neurodevelopmental outcome of each infant was predicted from an independent training sample. Cortical FA measurements from 52 parcellated cortical gyri formed the feature vectors of each subject and were used as the SVR predictor. Feature vectors for all subjects were concatenated (*Feature vectors* in Figure 1) to obtain the input data for SVR prediction models with linear kernel function (Figure 1). Each feature represented by FA measurement at each cortical gyrus was independently normalized across training data. Only training data was used to compute the normalization scaling parameters, which were then applied to the testing data. After predicted continuous cognitive or language scores were estimated by the prediction model, Pearson correlation coefficient (*r*) and mean absolute error (MAE) between the actual and predicted continuous score were computed to evaluate cognition or language prediction models. The normalized feature contribution weights (|*w*_*i*_|/ ∑|*w*_*i*_| with *i* indicating *i*th cortical gyrus) were calculated to represent contribution of all parcellated cortical gyri to the cognition or language prediction model. These normalized feature contribution weights of all parcellated cortical gyri in cognition or language prediction model were then mapped to the cortical surface to reveal heterogeneous regional contribution across entire cortex and distinguishable regional contribution distribution in a specific prediction model.

### Assessment of robustness of prediction

Permutation test was conducted to assess LOOCV prediction performance. Specifically, cognitive or language outcomes were randomly shuffled across subjects 1000 times. Prediction procedure was carried out with each set of randomized outcomes, generating null distributions. Pearson correlation was conducted for each set of randomized outcome. MAE between predicted and observed outcome from randomly shuffled distributions was also calculated. The p values of observed correlation coefficient (*r*) value in LOOCV prediction, calculated as the ratio of number of permutation tests with correlation coefficient greater than observed *r* value over number of all permutation tests, are the probability of observing the reported *r* values by chance. Similarly, the p values of MAE in LOOCV prediction, calculated as the ratio of number of permutation tests with MAE value lower than observed MAE value over number of all permutation tests, are the probability of observing the reported MAE by chance.

To investigate effect of cortical parcellation schemes on the cortical FA measures in prediction model, various cortical parcellation schemes, including 52 cortical regions from the neonate atlas labeling (Feng et al., 2019), 128, 256, 512 and 1024 randomly parcellated cortical regions with equal size (Zalesky et al., 2010) were tested. For each parcellation scheme, averaged value of skeletonized FA measurements in each cortical ROI was used as a feature in the SVR model to test prediction performance. To address a possible confounding factor of various neonate gestational ages at scan, we evaluated if the prediction performance of cortical FA measures remained high after controlling for ages at scan following the age correction methods described in the literature (Dukart et al., 2011). Specifically, age effect in the cortical FA of each gyrus (Ouyang et al., 2019) or each parcellated ROI was adjusted with linear regression between cortical FA and age. The cortical FA residuals in the linear regression model, considered as age adjusted cortical FA measures, were then used as features in SVR models for predicting the Bayley-III scores. To validate the capability of cortical FA in behavioral predictions, individual’s Bayley composite scores were categorized into normal (>85) and low scores (≤85). Cortical FA measures were used as features to classify each subject’s score into normal- or low-score groups using SVM algorithm with LOOCV. Classification accuracy and area under the receiver operating characteristic (ROC) curves were used to evaluate the performance of classification models.

### Bootstrap analysis for assessing reproducibility of top 10 cortical regions identified by LOOCV analysis

We used a bootstrap sampling approach to assess reproducibility of top 10 cortical regions where microstructural measures contributed most to predicting each outcome in LOOCV analysis. Specifically, we randomly selected 90% of the total 46 samples 1000 times. We then built cognition or language prediction model with each set of selected samples and identified top 10 cortical regions with highest contribution to the prediction of cognitive or language outcome. In each of 1000 bootstrap resamples, if any cortical region was identified as top 10 cortical regions contributing to predicting cognition outcome, the count for this specific cortical region was added by 1. After testing with 1000 resamples, a percentage of a certain cortical region was calculated as the total count for this region divided by 1000. In this way, a percentage map of all cortical gyri for predicting cognitive outcome can be created. The same procedure was repeated with 1000 bootstrap resamples for predicting language outcome. If the top 10 cortical regions where microstructural measures contributed most to predicting each outcome in LOOCV analysis (Figure 3) overlaps with the cortical regions with high percentage, it indicated that the top 10 cortical regions identified by LOOCV analysis were highly reproducible.

### Permutation tests to assess distinguishable regional contribution to predicting cognitive or language outcomes

To quantify the extent of distinction between the set of top 10 cortical regions in cognition prediction model and the set of top 10 cortical regions in language prediction model, we defined a nonoverlapping index as the number of nonoverlapped regions between these two sets divided by the number of regions in their union set. This nonoverlapping index ranges from 0 to 1, with 1 indicating completely distinctive sets of regions and 0 indicating completely same sets of regions. A permutation test was used to evaluate the statistical significance of the observed nonoverlapping index. The null hypothesis is that the observed nonoverlapping index from predicting two different outcomes is not different from a distribution of nonoverlapping index calculated from predicting same (cognitive or language) outcome. The null distribution of nonoverlapping indices was generated by calculating 2070 nonoverlapping indices with each corresponding to one of 1035 pairs of cognitive-cognitive outcome or one of 1035 pairs of language-language outcome using leave-one-out resamples. The p value of reported nonoverlapping index is the probability of observing the reported nonoverlapping index by chance, and was calculated as the number of permutations with higher index value than reported index divided by the number of total permutations. We also conducted a more strict permutation test by increasing variability of the resamples. Specifically, the bootstrap resamples used in the section of “bootstrap analysis for assessing reproducibility of top 10 cortical regions identified by LOOCV analysis” above was adopted to generate another null distribution of nonoverlapping indices and calculate the p value of observed nonoverlapping index by using the same procedures described above.

### Prediction of neurodevelopmental outcome with white matter FA as features

White matter microstructure quantified with dMRI was tested for predicting neurodevelopmental outcomes. White matter skeleton FA values at the core were measured to alleviate the partial volume effects (Figure 4a). White matter skeleton was further parcellated into 40 tracts with the tract labeling transformed from a neonate atlas (Feng et al., 2019). Details of tract-wise FA measurement at the white matter skeleton were described in our previous publication (Huang et al., 2012). White matter FA measurements of the 40 tracts were used to generate the feature vectors of each subject. Similar to the procedures of predicting neurodevelopmental outcomes with cortical FA feature vectors, these white matter FA feature vectors were the input of the SVR predictor with LOOCV for predicting neurodevelopmental outcome.

## Acknowledgements

This study is sponsored by NIH R01MH092535, R01MH092535-S1 and U54HD086984. We would like to thank Brittany C. Bennett at the Children’s Hospital of Philadelphia for her contribution to the schematic depiction.

## Figures and figure supplements

**Figure 1-figure supplement 1.**
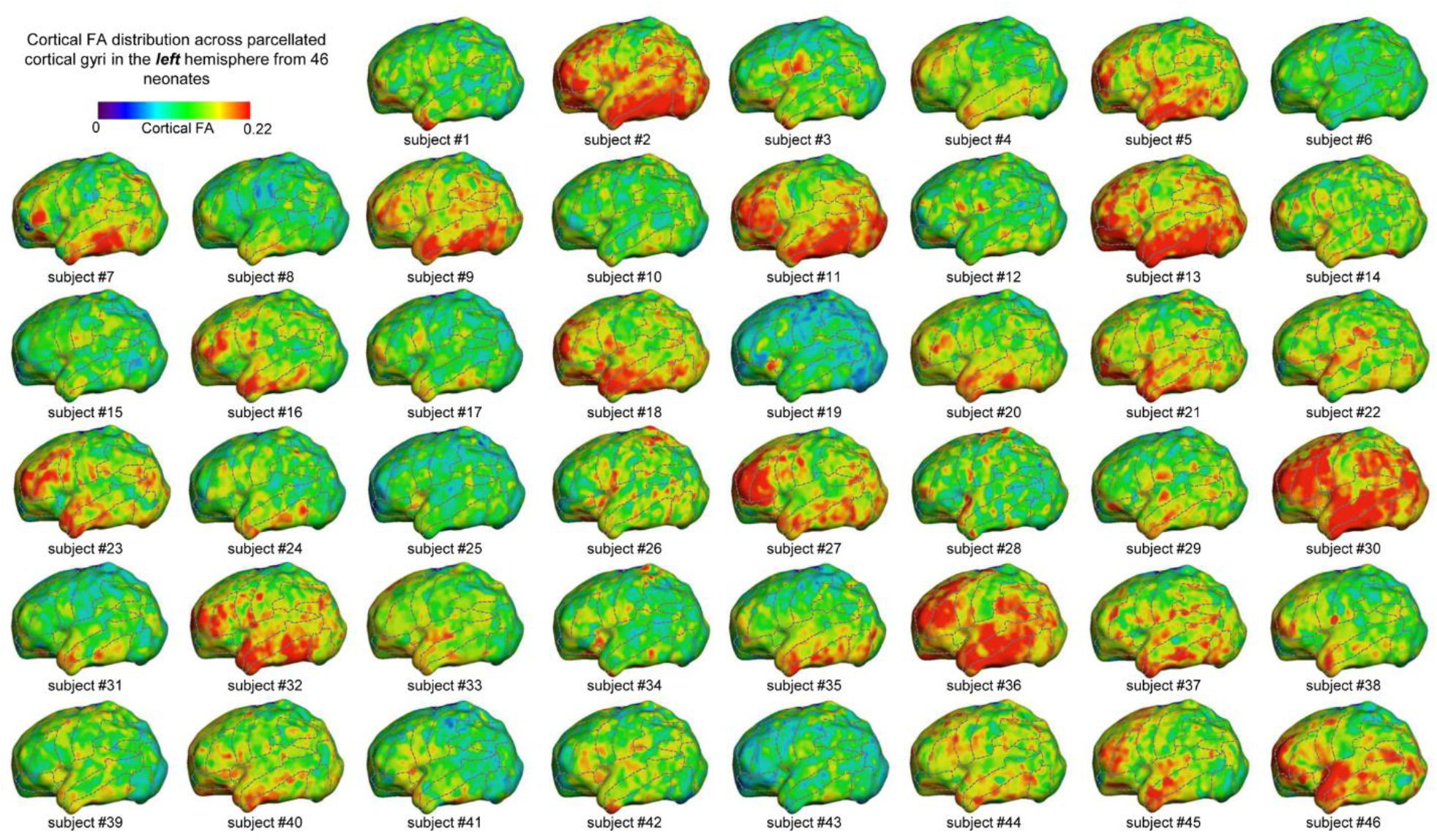
Cortical FA distribution across parcellated cortical gyri in the left hemisphere from 46 infants who also went through neurodevelopmental assessments with Bayley III at their 2-years of age. Colorbar on the upper left corner encodes the cortical FA values. The dashed lines demonstrate the boundaries among the parcellated cortical gyri.

**Figure 1-figure supplement 2.**
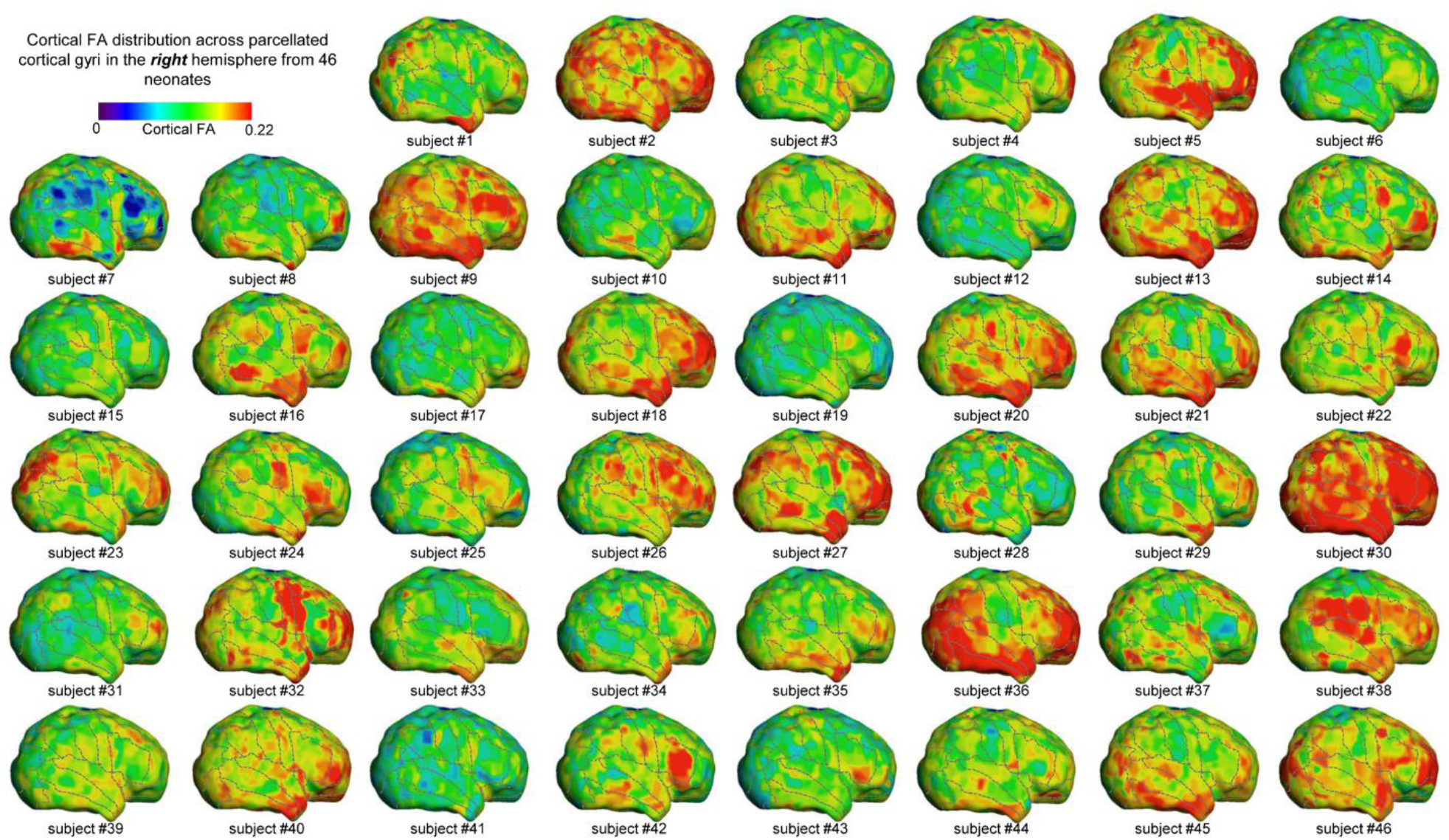
Cortical FA distribution across parcellated cortical gyri in the right hemisphere from 46 infants who also went through neurodevelopmental assessments with Bayley III at their 2-years of age. Colorbar on the upper left corner encodes the cortical FA values. The dashed lines demonstrate the boundaries among the parcellated cortical gyri.

**Figure 1-figure supplement 3.**
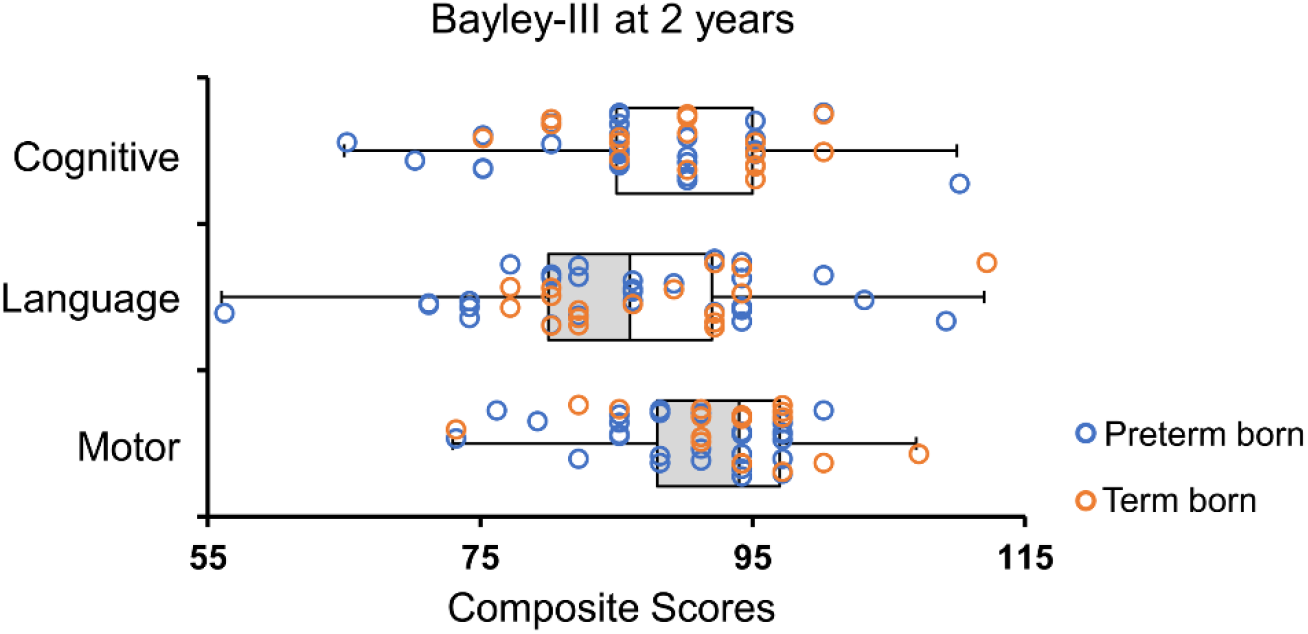
Distribution of the Bayley-III cognitive, language and motor composite scores of the studied population.

**Figure 2-figure supplement 1.**
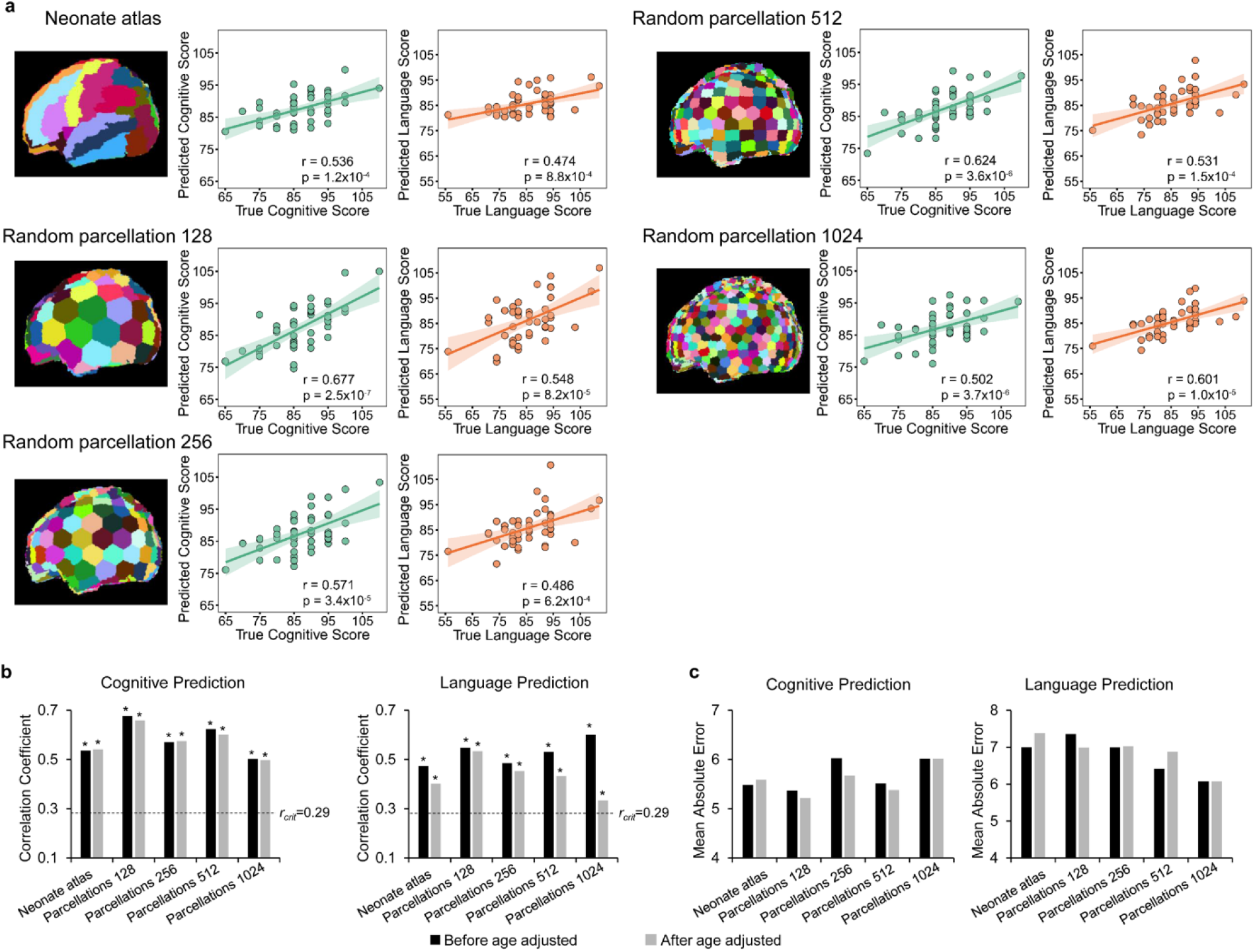
Evaluation of robustness of prediction models with different cortical region-of-interest (ROI) schemes, and age adjustments. **a**, Various cortical ROI schemes, including 52 cortical regions from the neonate atlas labeling, 128, 256, 512 and 1024 randomly parcellated cortical regions with equal size, were tested by using cortical FA measured at these ROIs as features in prediction models. The scatter plots show the linear regressions between actual scores and the predicted cognitive or language scores based on the cortical FA measures obtained with various ROI schemes. **b**, Significant correlations between the predicted and actual cognitive and language scores were found for all ROI schemes before age adjustment, and all ROI schemes after age adjustments. Dashed lines indicate critical r value corresponding to p=0.05. * in the panel indicates significant (p<0.05) correlation. **c**, Mean absolute error with all ROI schemes, and age adjustment are shown.

**Figure 2-figure supplement 2.**
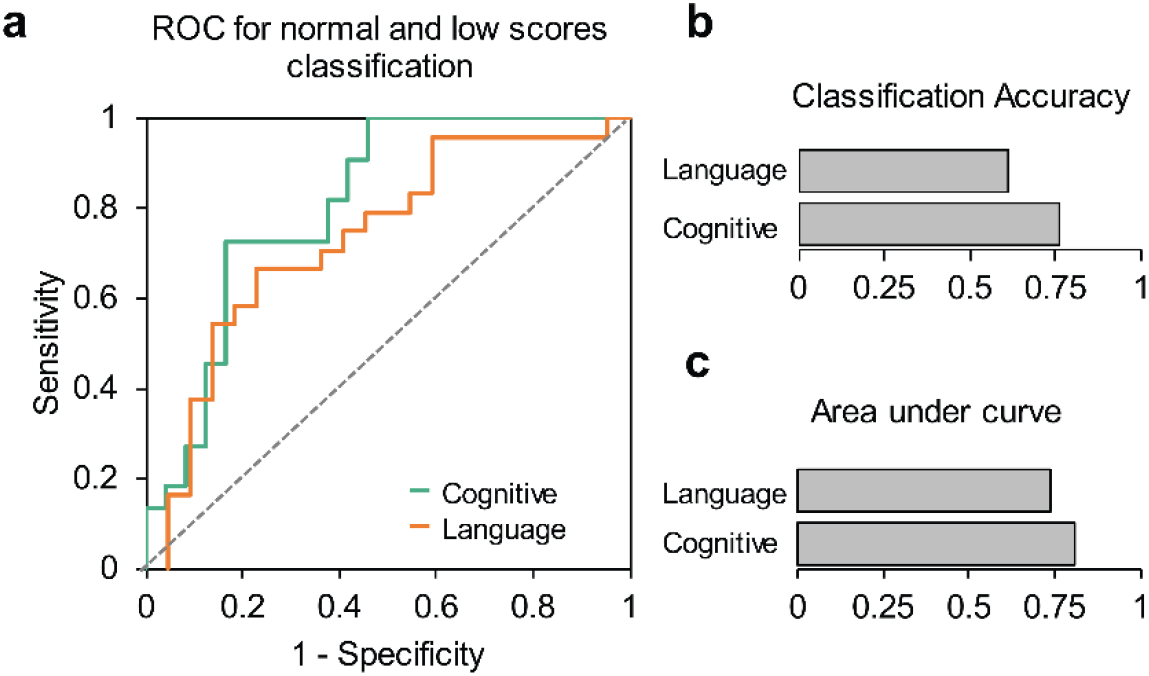
Evaluation of the prediction models. **a**, The receiver-operating-characteristic (ROC) plots show specificity and sensitivity with cortical microstructural measures used as features to classify normal (>85) versus low (≤ 85) cognitive or language scores. **b** and **c** show accuracy and area under ROC curve for language and cognitive classification model.

**Figure 3-figure supplement 1.**
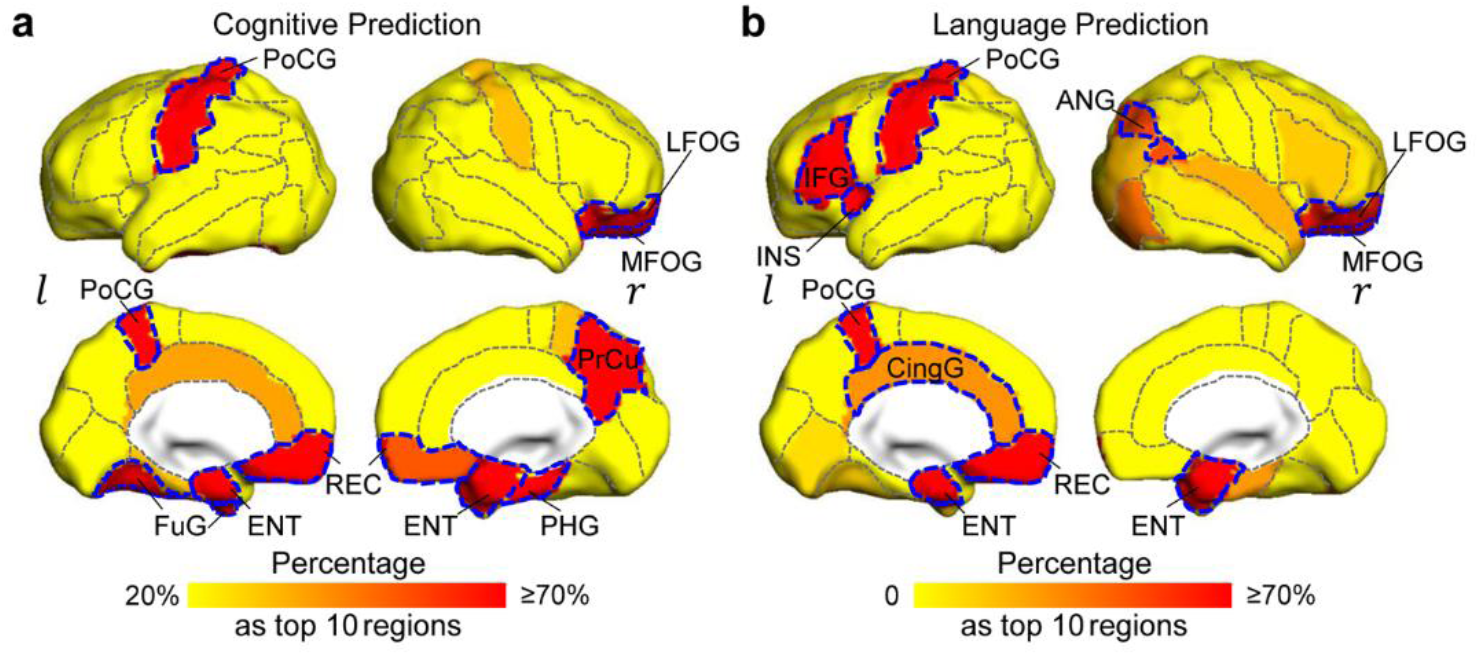
Percentage maps for evaluating reproducibility of top 10 cortical regions with highest feature contribution to predicting cognition (a) and language (b) outcomes from bootstrap analysis. Regions with high percentage (highlighted by red and brown) are consistent with the top 10 cortical regions (from Figure 3, highlighted by dashed blue contour), demonstrating high regional reproducibility for contributing to predicting a specific (cognitive or language) outcome.

